# Identification of host lncRNAs that impact Venezuelan equine encephalitis virus replication

**DOI:** 10.1101/2025.05.12.653438

**Authors:** Mahgol Behnia, Chunyan Ye, Kim Somfleth, Olufunmilola M. Oyebamiji, Kathryn J. Brayer, Yan Guo, Scott A. Ness, Ram Savan, Steven B. Bradfute

## Abstract

Venezuelan equine encephalitis virus (VEEV) causes encephalitis in humans and equids, and there are no vaccines or therapeutics available for humans. In recent years, non-coding RNAs have emerged as critical regulatory factors affecting different cellular pathways. Specifically, long non-coding RNAs (lncRNAs) have been identified as regulators of antiviral pathways during various viral infections; however, their role in regulating VEEV infection has not been assessed. Here we show differential expression of several lncRNAs in primary mouse target cells infected with a vaccine strain of VEEV (TC-83) but not a pathogenic strain (TrD). Among the differentially expressed genes (DEGs), suppressing lncRNA small nucleolar RNA host gene 15 (Snhg15) resulted in about a 7-fold increase in VEEV TC-83 replication in primary mouse astrocytes. Knockdown of Snhg15 during VEEV TC-83 infection resulted in the suppression of ten genes including Irf1, Junb, Atf3, Relb, Pim1, Hbegf, Ccl5, Ankrd33b, and H2-K2, all of which were also increased during TC-83 infection when the expression of Snhg15 increased in primary mouse astrocytes. Most of these genes are involved in antiviral responses. KEGG pathway analysis confirmed the suppression of both pattern recognition receptor and inflammatory pathways after in Snhg15 knockdown. These data are the first to identify lncRNA responses in encephalitic alphavirus infection and demonstrate important roles for these overlooked RNAs on VEEV infection.

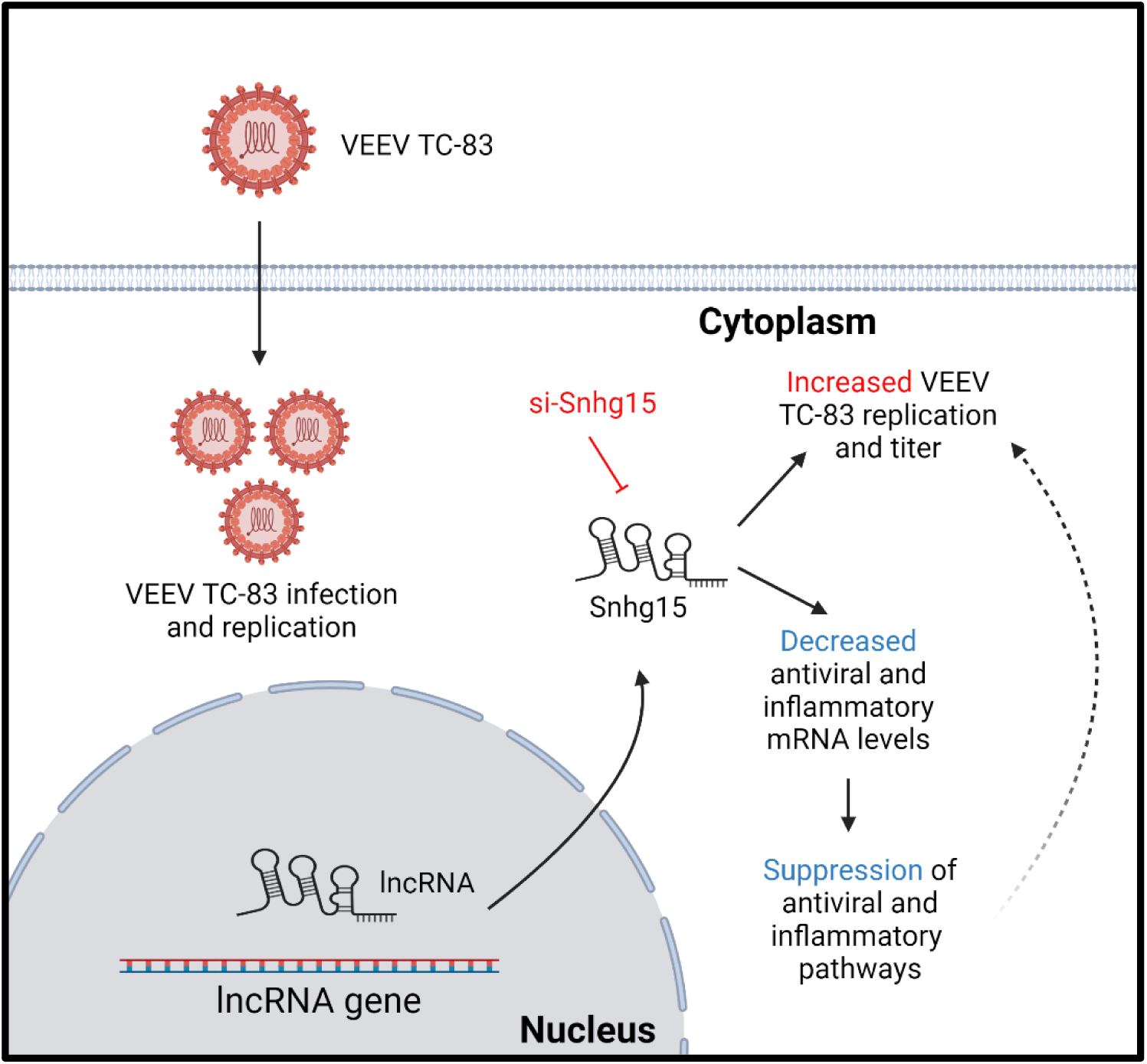

## Introduction

Venezuelan equine encephalitis virus (VEEV) can cause encephalitis in humans and equines and has been responsible for large encephalitic outbreaks in North, Central, and South America (Aguilar et al., 2011). VEEV is a positive-strand RNA virus from the alphavirus genus in the Togaviridae familyIn nature, both enzootic and epizootic strains of VEEV are transmitted through mosquito bites (Forrester et al., 2017). However, VEEV is also highly infectious through aerosols and has previously been developed as a biological weapon (Lundberg et al., 2017; Rusnak et al., 2018). These characteristics have led to the recognition of this virus as a potential biological terrorism agent, and it is classified as a Category B priority pathogen by the Centers for Disease Control and Prevention and the National Institutes of Health. VEEV Infection leads to abrupt onset of flu-like symptoms 2-6 day after infection, including fever, fatigue, headache, myalgia, and nausea in humans (Watts et al., 1998). Highly lethal in equids, VEEV disease is mostly self-limiting in humans, but in some cases causes encephalitis, resulting in a fatality rate of <1% in symptomatic patients. Despite the low fatality rate, 4-14% of patients develop encephalitis leading to permanent neurological sequela (Lundberg et al., 2017). Extensive efforts to produce a VEEV vaccine resulted in VEEV TC-83, which was generated by attenuation of a virulent select agent VEEV strain, VEEV Trinidad Donkey (TrD) (Pittman et al., 1996). However, VEEV TC-83 is only available to at-risk personnel and has questionable efficacy. The lack of an FDA-approved vaccine or therapeutic for human use underscores the necessity of investigating the VEEV-host interaction to identify new therapeutic targets.

Previous studies have reported the induction of type I and II interferon (IFN-I and IFN-II) and proinflammatory cytokines during VEEV infection in non-human primates, mouse models, and cell culture (Koterski et al., 2007; Sharma et al., 2008, 2011; Sharma & Knollmann-Ritschel, 2019; Sharma & Maheshwari, 2009; Valerol et al., 2008). However, other studies indicated that VEEV evades the host immune response by blocking expression of interferon stimulated genes (ISGs), inducing host transcriptional shutoff, and causing host translational shutoff, which are independently mediated by VEEV E2, capsid, and nonstructural protein 2 (nsP2), respectively (Bhalla et al., 2016; Eaton, 2021; Garmashova et al., 2007). Despite the ability of the viruses to evade innate antiviral responses, most viral infections can be cleared by the hosts suggesting the significance of additional host regulatory factors in modulation of the host innate antiviral pathways.

Long noncoding RNAs (lncRNAs) are a group of regulatory RNAs that are longer than >200 nucleotides in length and lack protein-coding potential (Mattick et al., 2023). Recent studies have identified lncRNAs as key regulators of signaling pathways that exert their regulatory effect through epigenetic, transcriptional, and post-transcriptional regulation of gene expression (Statello et al., 2021). lncRNAs regulate cellular processes like development, differentiation, and immune responses through regulation of gene expression (De la Fuente-Hernandez et al., 2022; Schmitz et al., 2016; Valadkhan & Gunawardane, 2016). Additionally, dysregulation of lncRNA expression is linked with different pathological conditions (Kaczynski et al., 2023; Shuai et al., 2019).

Notably, a growing body of studies report changes in cellular lncRNA expression during viral infections. These studies reveal that lncRNAs modulate innate antiviral responses through different mechanisms including regulation of pathogen recognition receptors (PRR) activation, activation of transcription factors, and antiviral gene expression (Chen et al., 2022; Vierbuchen & Fitzgerald, 2021; Xu et al., 2021). Moreover, altering the expression of differentially expressed lncRNAs regulates viral infection. lncRNA LINC02574, whose expression is induced upon influenza A virus (IAV) infection, is a positive regulator of the innate antiviral response, as suppression of this lncRNA led to higher IAV replication in A549 cells (Zhang et al., 2023). Conversely, siRNA-mediated disruption of lncRNA lncRHOXF1 in human trophectoderm progenitors during Sendai virus infection increased RIG-I and MDA-5 expression (Penkala et al., 2016). These studies show the importance of lncRNAs in modulation of viral infections, indicating that deeper understanding of lncRNA function during viral infection may lead to new cellular targets for treatment strategies.

In recent years, many studies have aimed to identify the function of lncRNAs during various viral infections. Despite this increasing effort, the lncRNA response to encephalitic alphaviruses and the regulatory impact of lncRNAs on their infection remains elusive. In this study, we investigated the lncRNA response to both wild-type (TrD) and vaccine strains (TC-83) of VEEV using mouse primary neurons and astrocytes, as VEEV-infected mice exhibit disease symptoms that closely resemble those observed in humans (Sharma & Knollmann-Ritschel, 2019)and provide a valuable system for investigating the antiviral responses to VEEV infection and disease. Importantly, utilizing primary cells known to be targets of VEEV can provide valuable and more biologically relevant information about a viral infection and the subsequent cellular response.

Here, our RNA sequencing analysis showed significant alterations in lncRNA expression following TC-83 but not TrD infection in primary mouse astrocytes and neurons in vitro. Subsequent experiments demonstrated increased TC-83 replication and titer in response to the suppression of three lncRNAs. Further investigation of the lncRNA Small nucleolar host gene 15 (Snhg15) indicated a substantial decrease in expression of antiviral genes after Snhg15 knockdown (KD) in primary mouse astrocytes infected with TC-83. Particularly, expression of rf1, Junb, Atf3, Relb, Pim1, Hbegf, Ccl5, Ankrd33b, and H2-K2 decreased in response to Snhg15 KD at all time points after infection. Further, Gene ontology and KEGG pathway analysis showed suppression of innate antiviral and inflammatory pathways in Snhg15 KD primary mouse astrocytes during VEEV TC-83 infection. Overall, our results suggest Snhg15 is a regulator of innate antiviral and inflammatory pathways during VEEV TC-83 infection.

## Results

### RNA sequencing reveals differential lncRNA expression in response to VEEV TC-83 but not VEEV TrD infection in primary mouse astrocytes and neurons

To explore the host lncRNA response to VEEV infection, we performed RNA sequencing in VEEV target cells, with or without infection with wild type (TrD) or live attenuated (TC-83) VEEV strains. Since astrocytes and neurons are primary VEEV target cells *in vivo*, we focused our experiments on these cells. Primary mouse neurons and astrocytes were infected with either VEEV TrD or VEEV TC-83 at multiplicity of infection (MOI) of 5 or left uninfected as controls for 16 and 24 hours. Total RNA obtained from the infected and uninfected control cells was fragmented before processing, as previously described (Behnia et al., 2022), to inactivate genomic viral RNA prior to RNA sequencing. Analyses of differentially expressed mRNAs revealed a significant increase in the expression of genes involved in antiviral signaling pathways across both cell types and in response to both infections. However, the number of differentially expressed genes (DEGs) and the extent of antiviral pathway activation were different based on viral strain. In both primary mouse astrocytes and neurons, the cellular mRNA response was more robust in TC-83 infection, evidenced by a strikingly higher number of genes being modulated in TC-83 infected cells compared to TrD infected cells (Figure 1A-B).

**Figure 1.**
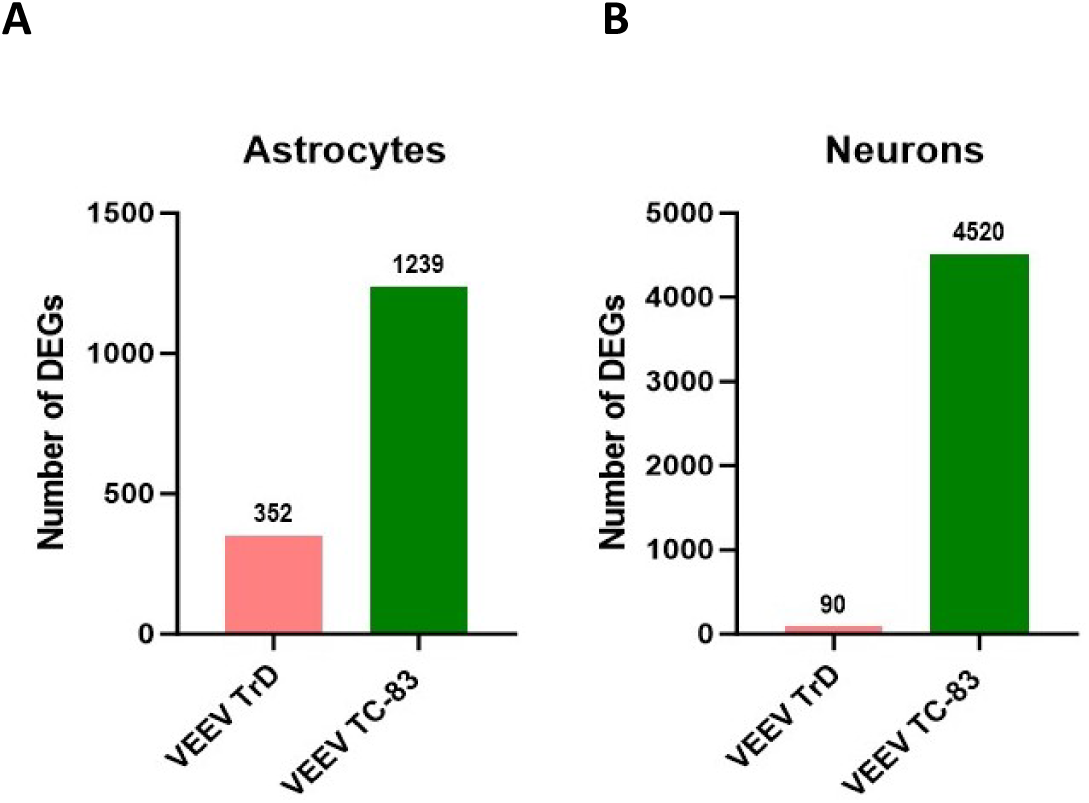
Cellular response to VEEV TrD vs VEEV TC-83 in different cell types in the CNS. Comparison of total number of differentially expressed mRNAs and lncRNAs in VEEV TrD-vs VEEV TC-83 infected primary mouse astrocytes **(A)** primary mouse neurons **(B)** at 24 hours. The genes with p.Adjust < 0.05 identified as significantly modulated in RNA-seq analysis.

Differential gene expression analyses indicated differential expression of a greater number of antiviral genes in TC-83 vs TrD-infected primary mouse astrocytes and neurons (Supplementary Figure1). These results show a vastly different cellular mRNA response to infections in wild-type versus attenuated VEEV. Further KEGG pathway analysis revealed activation of antiviral pathways in response to both strains of VEEV in primary mouse astrocytes and neurons (Supplementary Files 1 and 2). However, the number of enriched antiviral pathways was higher during TC-83 infection of these cells. These results suggest that the wild-type strain of VEEV efficiently evades the cellular antiviral responses in the central nervous system.

Notably, we observed a significant difference in the host lncRNA response to these infections. TC-83 infection led to substantial changes in the expression of cellular lncRNAs in primary mouse astrocytes and neurons. Particularly, 8 and 24 lncRNAs in primary mouse astrocytes and 22 and 78 lncRNAs in primary mouse neurons were differentially expressed at 16 and 24hpi, respectively (Supplementary Figure2A-B and Figure2A-B1). In contrast, TrD infection did not affect the expression of host lncRNAs in either cell type at these timepoints (Supplementary Figure 2C-D & Figure 2C-D).

**Figure 2.**
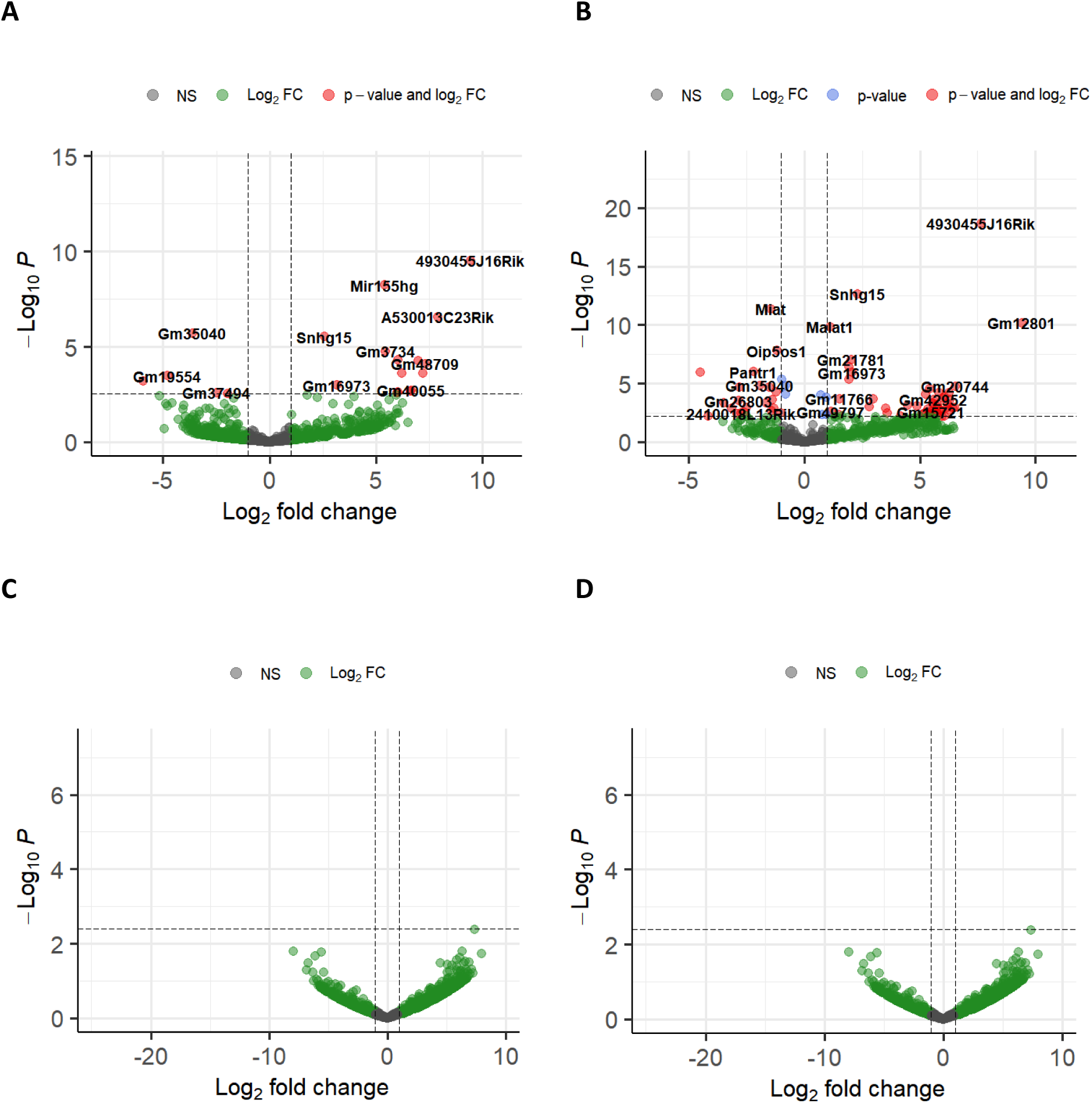
The host cellular lncRNA response to VEEV strains at 24hpi. **A-B)** Volcano plots show DE-lncRNAs in primary mouse astrocytes **(A)** and primary mouse neurons **(B)** infected with VEEV TC-83. **C-D)** Volcano plots show DE-lncRNAs in primary mouse astrocytes **(C)** and primary mouse neurons **(D)** infected with VEEV TrD. All the plots show the results from RNA-seq at 24h.p.i. The p-value threshold adjusted to show p.Adj value = 0.05.

Comparing the list of significantly altered lncRNAs in TC-83-infected primary mouse astrocytes and neurons revealed modulation of nine lncRNAs shared in both of these cells, with eight showing a consistent trend in both cell types at 24h.p.i. Interestingly, only two of these shared lncRNAs, Small nucleolar RNA host gene 15 (Snhg15) and Myocardial infarction associated transcript (Miat), have been previously characterized, while the function of the remaining lncRNAs are still unknown. These results demonstrate a distinct host lncRNA response to VEEV TC-83 infection in the cells of the central nervous system.

The vaccine strains of viruses elicit robust but safe immune responses to the viral infection. The robust mRNA and lncRNA responses to TC-83 infection but not pathogenic TrD observed in our study suggests activation of the pathways that are typically controlled or suppressed by wild-type VEEV. Given that lncRNAs can play key roles in regulating antiviral pathways, modulation of lncRNAs in response to TC-83 infection but not TrD infection suggests that these lncRNAs may serve as key regulators of host antiviral signaling pathways through modulating the viral infection.

### lncRNAs modulated in VEEV TC-83 infected primary mouse astrocytes regulate viral infection

To identify lncRNAs with anti-VEEV activity, we selected a subset of eight lncRNAs identified as differentially expressed in our RNA-seq analysis (Table 1). Among these, three lncRNAs, including Snhg15, Miat, and 4930455J16Rik, were significantly modulated in both TC-83-infected primary mouse astrocytes and neurons. The remaining five lncRNAs were differentially expressed only during TC-83 infection in primary mouse astrocytes. Although both astrocytes and neurons are vital VEEV target cells *in vivo*, because astrocytes are a place for VEEV replication and contribute to brain inflammation, we used these cells to investigate the potential regulatory function of modulated lncRNAs during VEEV infection.

**Table 1.** Differentially expressed lncRNAs selected for RNAi screening. p-adj and log2FC values at 24h.p.i.

| **lncRNA** | **Species** | **p-adjusted value** | **Log2 Fold change** |
| --- | --- | --- | --- |
| 4930455J16Rik | Mouse | 9.35E-08 | 9.427488 |
| Mir155hg | Human/Mouse | 9.22E-07 | 5.391059 |
| A530013C23Rik | Mouse | 2.84E-05 | 7.881687 |
| Snhg15 | Human/Mouse | 0.0002 | 2.561977 |
| Gm3734 | Mouse | 0.0009 | 5.455434 |
| Miat | Human/Mouse | 0.02 | -5.92181 |
| 3110039M20Rik | Mouse | 0.04 | 6.035248 |
| Mir9-3hg | Human/Mouse | 0.05 | -1.96377 |

To investigate the potential regulation of VEEV infection by differentially expressed lncRNAs, we began by validating the expression change in the selected lncRNAs during TC-83 infection using RT-qPCR. To this end, we infected primary mouse astrocytes with TC-83 (MOI 5) or left them uninfected as controls. The cells were lysed at 16 and 24hp.i. and total RNA extracted from them were subjected to TaqMan assays targeting the lncRNA of interest and GAPDH as an internal control. These validations were performed in five biological replicates. Among the eight selected lncRNAs, Snhg15, Mir9-3hg, Miat, A530013C23Rik, and 3110039M20Rik showed expression changes consistent with our RNA-seq results in at least three biological replicates at both timepoints post-infection (Figure 3).

**Figure 3.**
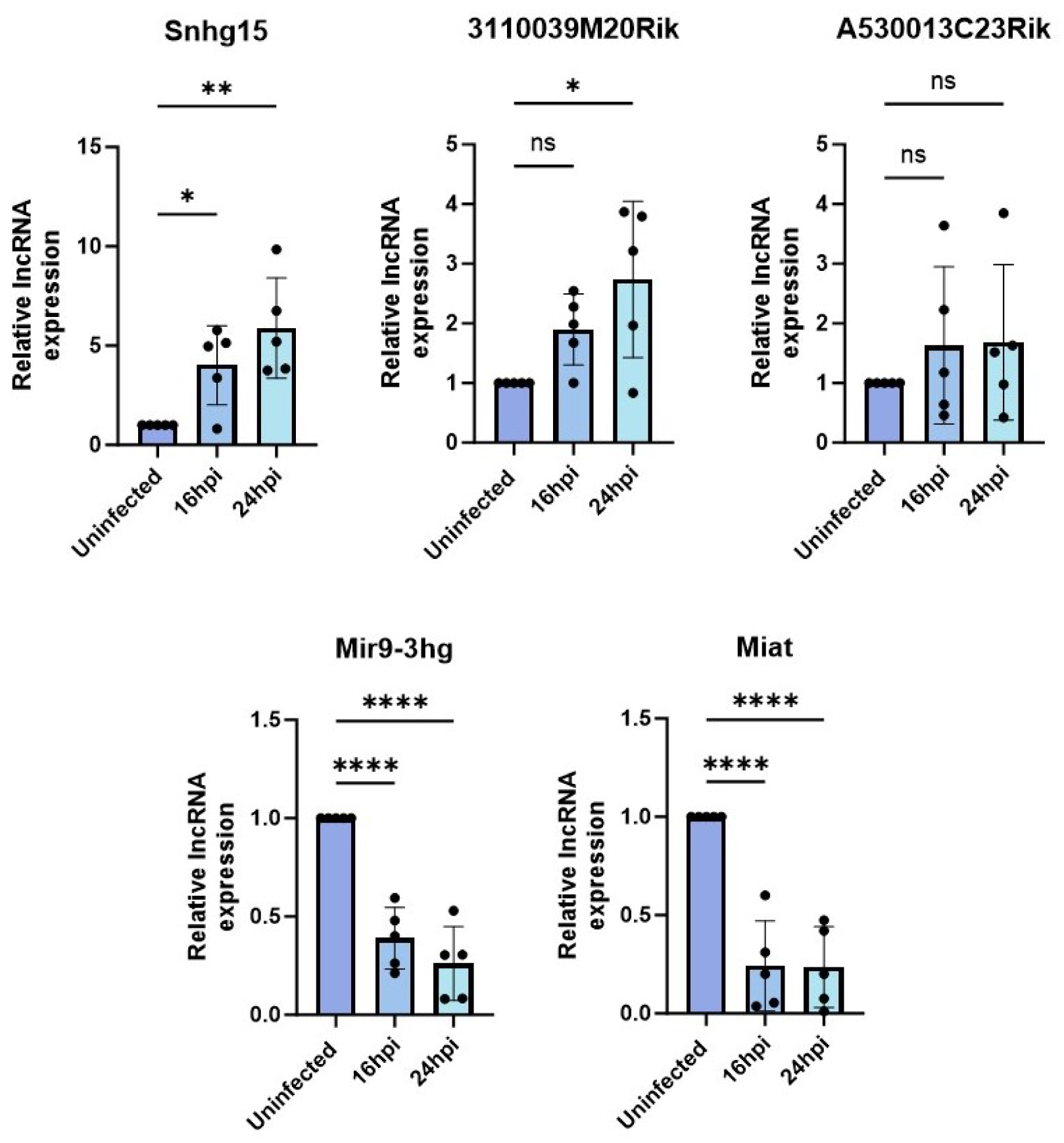
Validation of the lncRNA expression using RT-qPCR. Expression changes in five lncRNAs were successfully validated in VEEV TC83 infected (MOI 5) primary mouse astrocytes at 16 and 24hpi using TaqMan assays. The expression of each lncRNA measured relative to GAPDH and normalized to lncRNA expression level in the uninfected control. Each dot represents a biological replicate. One-way ANOVA followed by multiple comparison used to measure the statistical significance.

Next, we performed an RNAi screen to investigate the potential regulatory impact of these validated lncRNAs on TC-83 replication and titer. For this purpose, we first confirmed lncRNA suppression using RT-qPCR. To this end, we transfected primary mouse astrocytes with siRNAs targeting the lncRNA of interest and then infected them with TC-83 (MOI 5). The lncRNA expression in this group was compared to cells transfected with non-targeting siRNA (to measure basal lncRNA expression) and cells transfected with non-targeting siRNA followed by TC-83 infection (to measure lncRNA expression changes during infection). Among the five validated lncRNAs, expressions of four (Snhg15, A530013C23Rik, Mir9-3hg, and Miat) were successfully suppressed using siRNAs in TC-83 infected primary mouse astrocytes (Figure 4).

**Figure 4.**
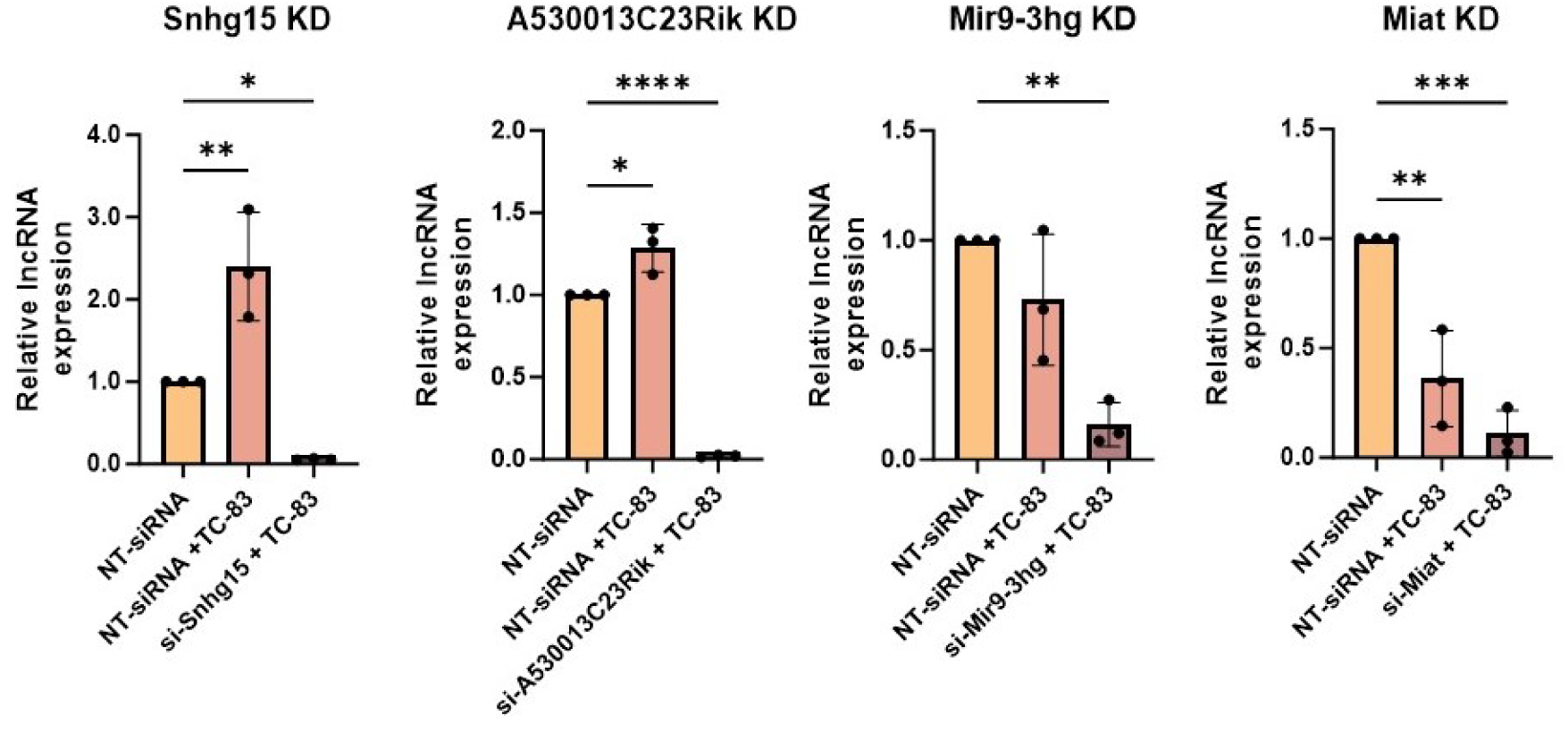
Four lncRNAs were suppressed using siRNAs in TC-83-infected primary mouse astrocytes. lncRNA suppression using siRNA was validated using RT-qPCR. The cells were transfected with either NT-siRNAs or si-lncRNA followed by infection with TC-83 at MOI 5 for A530013C23Rik KD, Mir9-3hg KD, Miat KD and MOI 0.5 for Snhg15 KD cells. lncRNA expressions were measure using TaqMan assays at 24hpi. The lncRNA expression calculated relative to GAPDH and normalized to the lncRNA expression level in uninfected cells. Each symbol represents a biological replicate (n=3). The statistical significance of lncRNA expression change tested using one-way ANOVA with multiple comparison.

To evaluate the regulatory impact of these three lncRNAs on VEEV replication, we performed TaqMan assays targeting VEEV capsid RNA in lncRNA-suppressed cells and controls. Primary mouse astrocytes transfected with a pool of four non-targeting siRNA infected with VEEV TC-83 (MOI 5) for 24 hours served as a control for viral replication. Changes in the number of VEEV TC-83 genome were quantified by generating a standard curve using a plasmid containing the complete VEEV TC-83 sequence in our RT-qPCR assays. Our results showed that the suppression of all four lncRNAs increased VEEV RNA copy number, with Snhg15 knockdown leading to 7-fold increase (Figure 5A). To further investigate the effect of lncRNA suppression on VEEV titer, we collected the supernatants from lncRNA knockdown and control cells and performed plaque assay. Consistent with the RT-qPCR results, suppression of all four lncRNAs increased VEEV TC-83 titer, with lncRNAs A530013C23Rik and Snhg15 suppression resulting in the highest increase (Figure 5B). These findings suggest the selected lncRNAs are regulators of VEEV replication. Among the four lncRNAs studied, suppression of Snhg15 had a pronounced effect on both viral RNA copy number and the number of infectious virions. Therefore, we selected Snhg15 for further functional analysis during VEEV infection.

**Figure 5.**
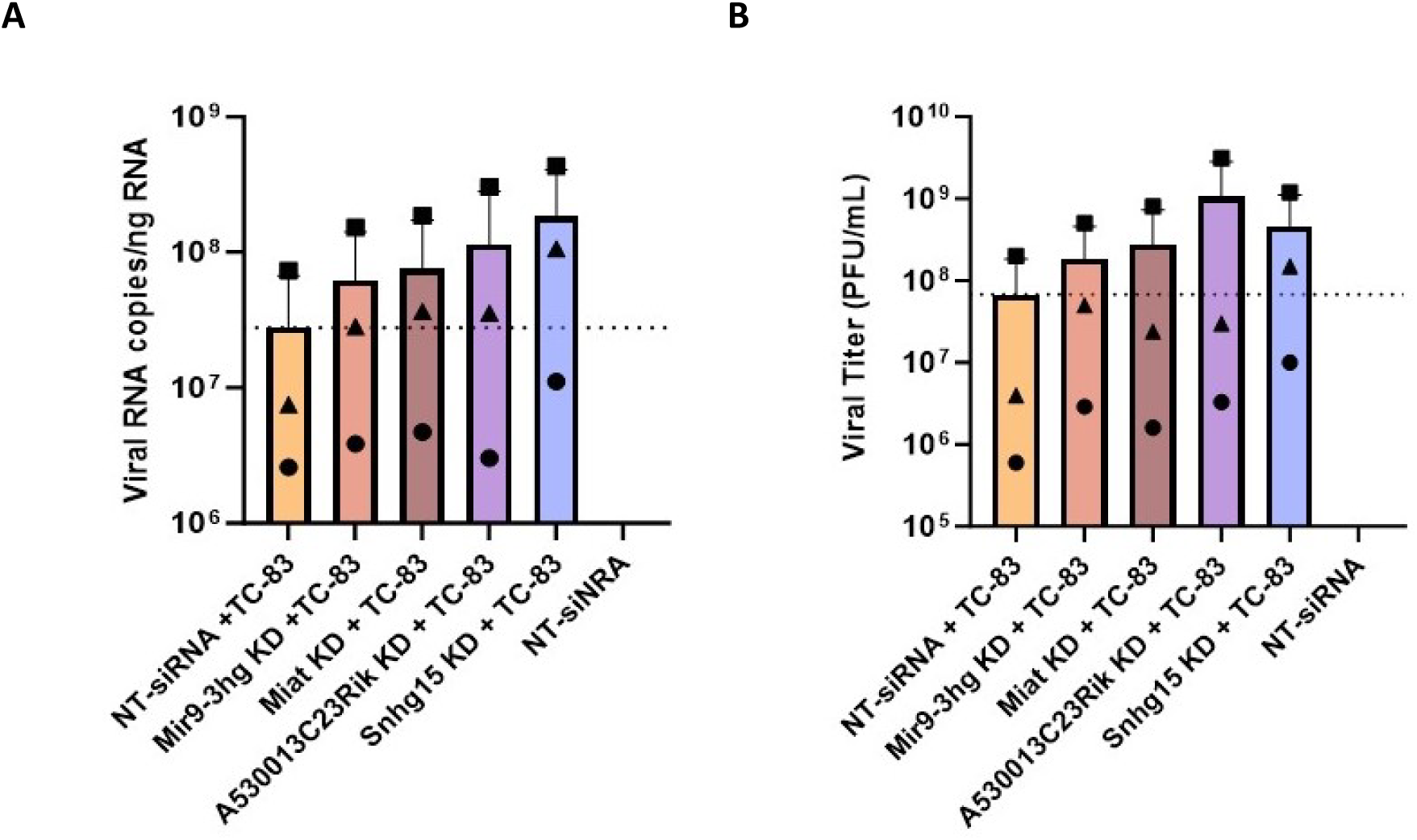
Suppression of three lncRNA changed VEEV TC-83 replication and titer. **A)** The effect of lncRNA KD on VEEV TC-83 RNA copy number measured using TaqMan assay targeting VEEV capsid. Capsid copy numbers measured using a standard curve generated based on the VEEV TC-83 plasmid and divided by ng of RNA used in each reaction. **B)** lncRNA suppression increased VEEV titer. Supernatants from lncRNA suppressed cells which are infected with VEEV TC-83 (MOI 5) for 24 hours used to perform plaque assays. Plaques were counted at 48hpi and PFU/mL calculated for all samples. Shapes correspond to the same biological sample across conditions.

### lncRNA Snhg15 may regulate VEEV TC-83 replication by modulating antiviral response to infection

After validating the effect of Snhg15 suppression on VEEV replication and titer, we next measured the impact of Snhg15 knockdown on antiviral gene expression during TC-83 infection. Using RNA sequencing, we analyzed gene expression in primary mouse astrocytes with and without Snhg15 suppression at 8, 16, and 24 hours after TC-83 infection. Control cells were transfected with a pool of four non-targeting siRNAs before infection with TC-83 to account for any transfection and infection effects on gene expression.

Our differential gene expression analysis revealed significant alterations (pAdj < 0.05, log2FC > 1) in the expression of 298, 197, and 352 host genes between Snhg15KD and control cells at 8-, 16-, and 24-hours post-infection, respectively. Notably, the expression of 124 genes were changed across all time points after TC-83 infection in Snhg15KD cells (Figure 6A). Interestingly, many of these genes encoded proteins involved in antiviral signaling pathways, suggesting Snhg15 as a regulator of antiviral pathways (Figure 6B-D). To identify genes directly affected by Snhg15 expression, we compared the expression changes in these 124 genes with their expression changes observed during TC-83 infection, which was obtained from our primary RNA-seq data. This analysis revealed that a subset of ten genes including Activating transcription factor 3 (Atf3), Interferon regulatory factor 1 (Irf1), Junb, Relb, Pim1, Histocompatibility 2, K region locus 2 (H2-K2), C-C motif chemokine ligand 5 (Ccl5), Heparin binding EGF like growth factor like (Hbegf), NFKB inhibitor epsilon (Nfkbie) and ankyrin repeat domain 33B (Ankrd33b), exhibited a significant increase in expression during TC-83 infection, coinciding with increased Snhg15 expression. Importantly, the expression of these genes was suppressed during Snhg15KD in the presence of TC-83 infection. Notably, most of these genes are involved in cellular antiviral responses. For example, Irf1, Relb, and Atf3 are transcription factors that have been previously studied for their roles in regulating antiviral responses (Badu & Pager, 2023; Panda et al., 2019; Saha et al., 2020).

**Figure 6.**
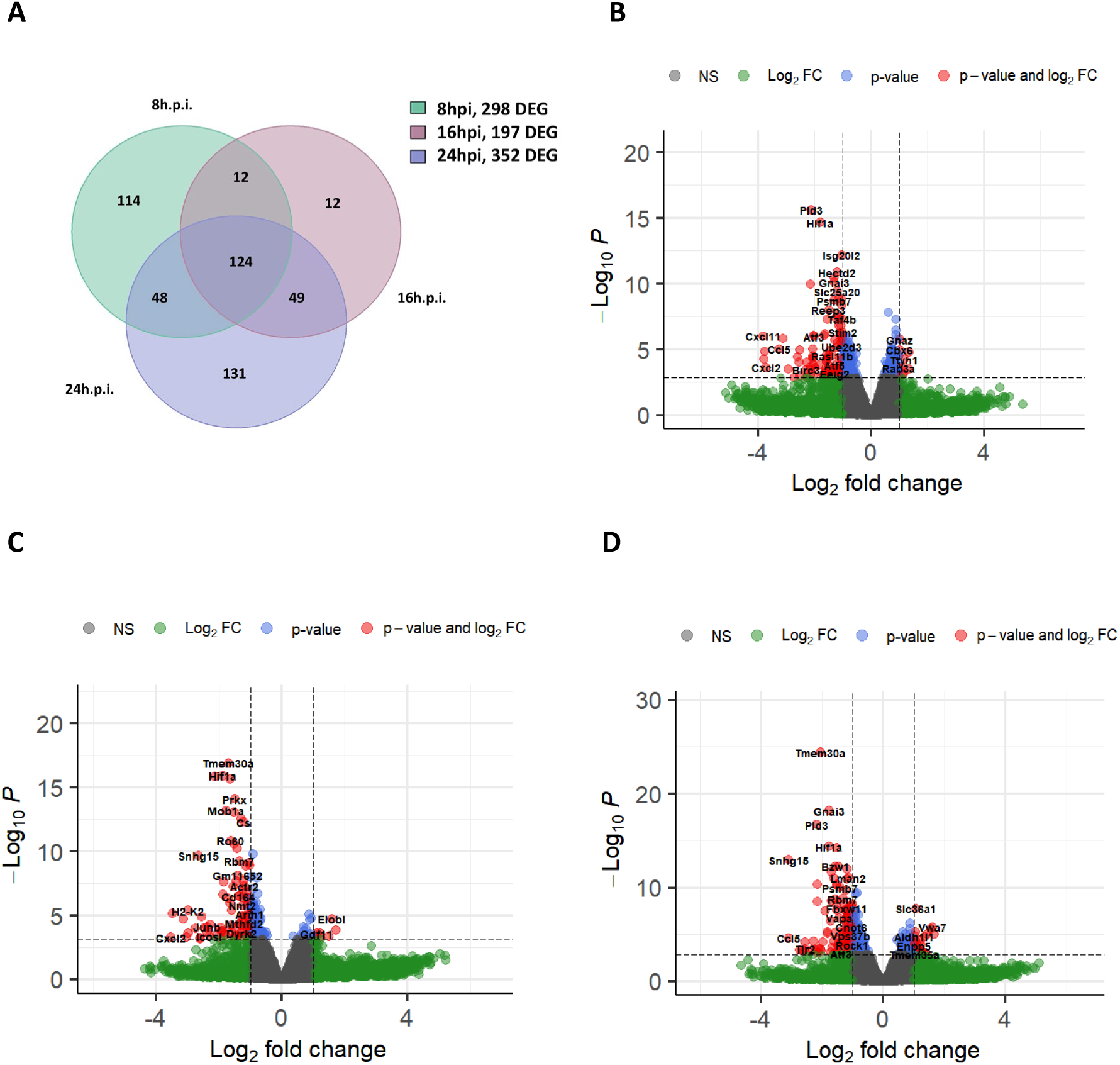
Snhg15 suppression decreased expression of genes involved in inflammatory response to viral infection. **A)** The Venn diagram shows the number of differentially expressed genes in Snhg15 suppressed and NT-siRNA treated primary mouse astrocytes, at 8, 16, and 24hpi with VEEV TC-83 (MOI 5). **B-D)** volcano plots of differentially expressed genes in Snhg15KD vs non-targeting siRNA treated primary mouse astrocytes at different timepoints post infection with VEEV TC-83 (MOI 5) at 8hpi**(B)**, 16hpi**(C)**, and 24hpi**(D)**.

Additionally, decreased transcription of Tlr2, a pattern recognition receptor involved in the innate antiviral response, was observed in Snhg15KD cells at 16- and 24hpi. The chemokines Cxcl1 and Cxcl2, which acts as chemoattractant for various immune cells during viral infection, were substantially decreased at 8- and 16-hours post-infection in Snhg15 suppressed cells. We investigated our RNA-seq data from TC-83-infected primary mouse astrocytes to examine the expression levels of Tlr2, Cxcl1, and Cxcl2. We observed that these genes were upregulated during infection, coinciding with the increased expression of Snhg15 in response to TC-83 infection. This observation might suggest a more direct Snhg15 regulatory impact on these genes. Overall, the suppression of genes involved in antiviral responses, along with increased TC-83 replication and titer observed in Snhg15KD cells, suggests that Snhg15 is a critical regulator of the antiviral response to VEEV infection.

To further investigate the potential regulatory impact of Snhg15 on the antiviral cellular response during VEEV infection, we performed KEGG pathway analyses. As expected, these analyses revealed the downregulation of several antiviral pathways in Snhg15 knockdown (KD) cells during TC-83 infection. Notably, key innate antiviral pathways, such as the JAK-STAT signaling pathway which is responsible for the production of type I interferons and their downstream signaling, were suppressed in Snhg15 KD cells at earlier time-points of infection. This finding coincided with a non-statistically significant decrease in the transcript level of interferon β (IFN-β) which was observed in Snhg15 KD cells (6-fold decrease at 8 hpi, 3.8-fold at 16 hpi, and 2.7-fold at 24 hpi). Additionally, a comparison of the all enriched pathways in Snhg15 KD cells at different time points post-infection showed the suppression of pathways involved in viral pathogen recognition such as TLR, RIG-I like receptor, NOD like receptor, and c-type lectin receptor signaling pathways. Moreover, Snhg15 suppression was associated with downregulation of inflammatory pathways such as NF-kB, TNF, and IL-17 signaling pathways, all of which are involved in the host antiviral response (Figure 7A-C). These findings suggest that Snhg15 serves as a positive regulator of the cellular antiviral responses during VEEV infection.

**Figure 7.**
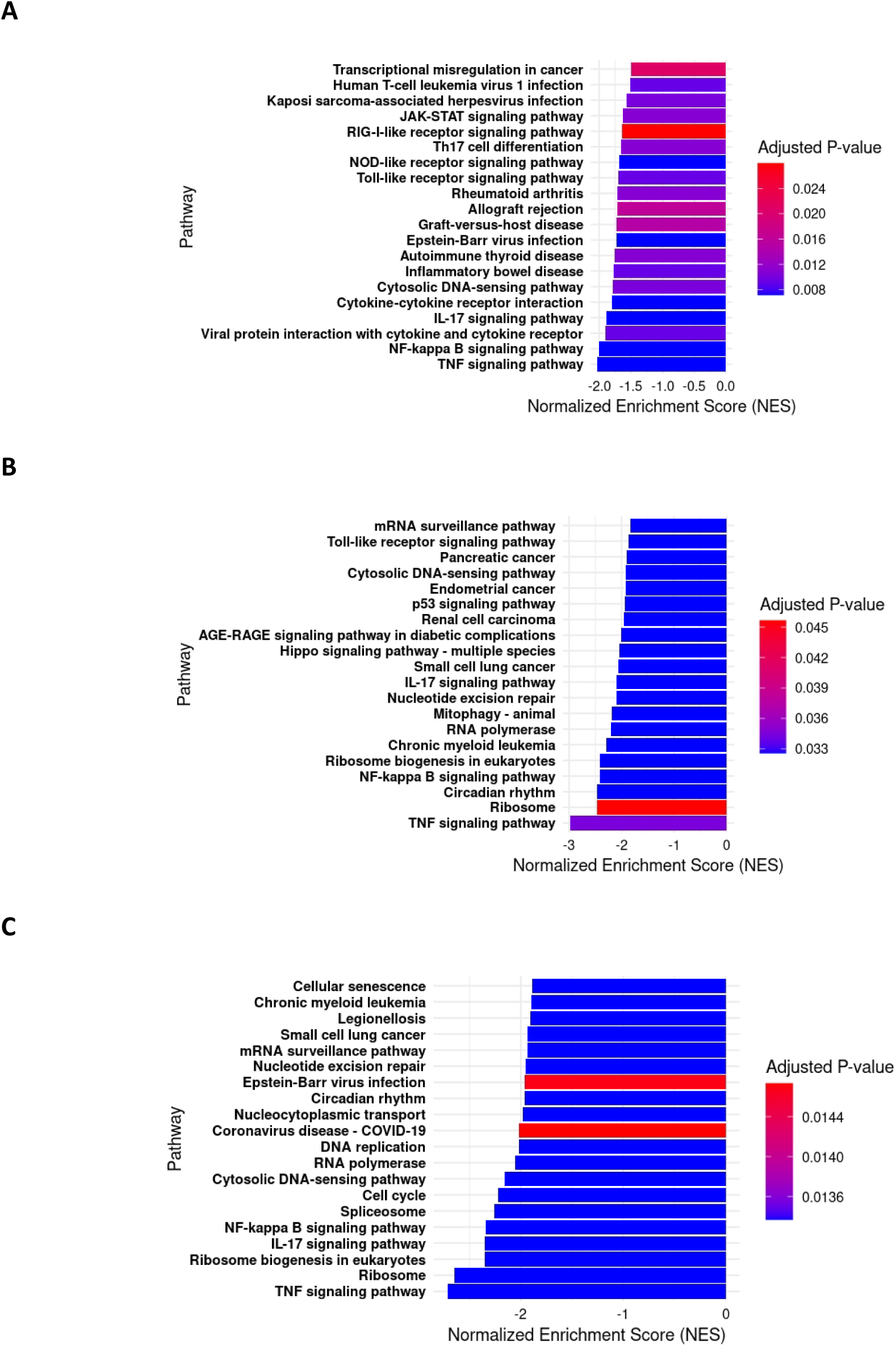
Snhg15 suppression antiviral pathways. **A-C)** Barplots of top20 negatively enriched KEGG pathways in Snhg15KD primary mouse astrocytes at 8hpi **(A)**, 16hpi**(B)**, and 24hpi **(C)** with VEEV TC-83 (MOI 5). The p.Adj cut off value for the KEGG pathway analysis is <0.05.

## Discussion

Modulation of cellular antiviral responses by lncRNAs has been reported during some viral infections (Ginn et al., 2021). However, the lncRNA response to encephalitic response to VEEV infection and introduce the regulatory impact of lncRNAs on this viral infection. Our RNA-seq analysis showed significant modulation of several lncRNAs in response to TC-83 infection in primary mouse astrocytes and neurons. However, this modulation was not observed in TrD infection of these cells (Supplementary Figure 2 and Figure 2).

The observed differences in lncRNA responses to various strains of VEEV in astrocytes and neurons may be due to differences in cellular antiviral responses to these infections. We identified 1,088 protein-coding genes whose expression specifically changed in response to TC-83 infection in primary mouse astrocytes at 24hpi, many of which are essential for antiviral responses. In contrast, only a few genes involved in antiviral responses were upregulated in TrD-infected astrocytes, all of which were also increased during TC-83 infection. These findings align with previous reports indicating a more robust antiviral response to partially neurovirulent VEEV (VEEV 3034) compared to neurovirulent VEEV (VEEV 3000) infection in mice (Gupta et al., 2017). The increased expression of genes involved in antiviral and inflammatory responses in TC-83-infected astrocytes suggests the activation of antiviral pathways in response to this strain of the virus that are potentially evaded by TrD. KEGG pathway analyses confirmed the activation of a greater number of antiviral signaling pathways in TC-83-infected astrocytes and neurons compared to TrD-infected cells, some of which exclusively enriched during infection with the vaccine strain (Supplementary Files 1 and 2). Particularly, several innate antiviral pathways such as RIG-I like receptor signaling, C-type lectin receptor signaling, and MAPK signaling pathways, were positively enriched in TC-83 infection of astrocytes, but not in TrD infection of these cells (Supplementary File 1). Similarly, JAK-STAT signaling pathway, NOD like receptor signaling pathway, and NF-kB receptor signaling pathway were identified as positively enriched in TC-83-infected, but not TrD-infected primary mouse neurons (Supplementary File 2).

We demonstrated that suppression of four lncRNAs, including Snhg15, Mir9-3hg, A530013C23Rik, and Miat, regulate VEEV viral titers, as measured by capsid RNA levels in cells as well as plaque-forming virus in supernatants (Figure 5). Using RNA-seq and differential gene expression analysis, we identified a subset of ten genes (Atf3, Irf1, Junb, Relb, Nfkbie, Ccl5, Pim1, H2-K2, Hbegf, and Ankrd33b) whose expression was positively correlated with Snhg15 expression in Snhg15 KD and normal Snhg15 expressing TC-83 infected primary mouse astrocytes at all timepoints after infection (Table 2). Most of these genes are known regulators of antiviral and inflammatory responses, showing that Snhg15 is a regulator of antiviral and inflammatory responses during VEEV infection.

**Table 2.** Genes whose expression positively correlated with Snhg15 expression and their functions.

| Gene | Function |
| --- | --- |
| Irf1 | Regulation of antiviral and inflammatory responses through induction of IFNβ, ISGs, TLR2, and TLR3 |
| Nfkbie | Regulation of inflammatory response through inhibition of NF-κB signaling pathway |
| Relb | NF-κB transcription factor involved in Regulation of NF-κB induced genes |
| Ccl5 | Chemokine involved in recruitment of leukocytes to the site of inflammation. |
| Junb | AP1 transcription factor involved in cellular response to stress |
| Atf3 | Stress-induced transcription factor involved in regulation of antiviral gene expression e.g. STAT1 |
| Pim1 | Serin-Threonine kinase involved in suppression of interferon signaling and ISG production |
| Hbegf | Involved in Influenza A entry |
| Ankrd33b | Function unknown |
| H2-K2 | Function unknown |

Snhg15 suppression decreased the expression of interferon regulatory factor 1 (Irf1), a transcription factor that plays a key role in the antiviral immune response by inducing early Interferon β (IFNβ) expression and the production of ISGs (Nair et al., 2017; H. Zhou et al., 2022). Additionally, IRF1 controls expression of Toll like receptor 2 and 3 (TLR2 and TLR3) (Panda et al., 2019), through which it can regulate inflammatory response to viral infection. Depletion of IRF1 has reported to reduce the protective efficacy of vaccination against virulent VEEV in a mice model of the disease (Grieder & Vogel, 1999), suggesting the importance of this transcription factor in host response to VEEV infection potentially though modulation of IFN-I and inflammatory response. Although we observed only a small decrease (from 6-fold to 3-fold) in IFN-β transcript levels in Snhg15KD cells throughout the entire infection period, TLR2 expression decreased significantly at 16 and 24hp.i. Notably, the regulatory effects of Snhg15 on IRF gene expression have not been studied; however, our RNA-seq data suggest that Snhg15 may affect VEEV infection by modulating Irf1 expression, subsequently affecting the expression of IFN-β and Tlr2 post-infection.

TLR2 is a pattern recognition receptor that can recognize viral pathogen associated molecular patterns (PAMP). TLR2 signaling activates NF-kB, MAPK, and IRF1/STAT1 pathways. Activation of these pathways lead to expression of downstream genes, including IFN-I and pro-inflammatory mediators. Snhg15 suppression resulted in a significant decrease in the transcript levels of key components of the NF-kB signaling pathway, such as Nfkbie (an inhibitor of NF-kB) and Relb (a Nf-kB transcription factor), across all time points post-infection. Furthermore, Snhg15 suppression led to decreased expression of Cxcl1 and Cxcl2 at 8- and 16hpi, as well as Ccl5 at all time points in TC-83-infected primary mouse astrocytes. Notably, the Nfkbie, Relb, Cxcl1, Cxcl2, and Ccl5 expressions are induced by NF-kB transcription factors. Notably, Snhg15 has been shown to interact directly with NF-kB signaling pathway to modulate the expression of inflammatory cytokines, such as IL6, IL1β, and TNFα (Sun et al., 2022, p. 202). Therefore, the decreased expression of these genes could point to reduced activation of NF-kB signaling in response to Snhg15 suppression.

Besides induction by NF-kB transcription factors, the expression of inflammatory mediators can be induced by Activator protein-1 (AP1) transcription factors in response to MAPK signaling activation (Korbecki et al., 2022; Okamura et al., 2020; Wickremasinghe et al., 2004). Interestingly, Snhg15 suppression decreased transcript levels of Junb, a key transcription factor in the AP1 family. Considering AP1 transcription factor’s involvement in the inflammatory response, the decreased transcript levels of Junb in response to Snhg15 suppression may affect the inflammatory response to VEEV infection.

Snhg15 suppression significantly reduced the mRNA levels of Activating Transcription Factor 3 (Atf3) and Proto-Oncogene Serine/Threonine Kinase (Pim1) in primary mouse astrocytes following TC-83 infection. ATF3 is a stress induced transcription factor which modulates genes involved in different cellular processes including inflammation and antiviral responses (Badu & Pager, 2023; Hai et al., 2018). ATF3 enhanced the expression of the antiviral genes such as STAT1, thereby increased the expression of other innate antiviral genes and controlled Zika virus infection (Badu & Pager, 2023). However, siRNA depletion of ATF3 decreased TC-83 infection to about 40%, suggesting ATF3 as a proviral factor during VEEV infection (Yao et al., 2021). Similarly, Pim1 promoted Zika infection by inhibiting STAT1 and STAT2 phosphorylation, thereby reducing interferon signaling and ISG production (F. Zhou et al., 2021). These findings indicate that the functions of ATF3 and PIM1 warrant further investigation into the effects of Snhg15 modulation on these factors during VEEV infection.

In addition to regulating innate antiviral response genes, Snhg15 suppression decreased the expression of Hbegf, Ankrd33b, and H2-K2 in VEEV TC-83-infected primary mouse astrocytes. Heparin-binding EGF-like growth factor (HBEGF) has been shown to reduce influenza A virus (IAV) titers in A549 cells (Lai et al., 2020). The functions of ankyrin repeat domain 33B (Ankrd33b) and histocompatibility 2, K region locus 2 (H2-K2) are not known. Whether Snhg15 modulates VEEV infection through modulation of these genes warrants further investigation.

Overall, our gene expression analysis in TC-83-infected primary mouse astrocytes revealed modulation of genes involved in the innate antiviral response and the proinflammatory response as a result of Snhg15 suppression. The decreased expression of Tlr2, Relb, Nfkbie, Cxcl1, Cxcl2, Ccl5, Pim1, and Junb suggest decreased enrichment of TLR and NF-kB signaling pathways. KEGG pathway analysis confirmed the negative enrichment of inflammatory pathways at all time points post-infection in Snhg15 knockdown cells, with TNF-α, NF-kB, and IL-17 signaling among the top ten suppressed pathways (Figure7A-C). Additionally, KEGG pathway analysis showed negative enrichment of RIG-I like receptor signaling, NOD-like receptor and C-type lectin receptor signaling in response to Snhg15 suppression, all of which are involved in host antiviral responses. Furthermore, signaling pathways that are involved in host defense against other viral infections like pathways activated during Coronavirus disease, Epstein-Barr virus infection, Human T-cell leukemia virus 1, Kaposi sarcoma associated herpes virus infection, Hepatitis B virus, Hepatitis C virus, Human cytomegalovirus infection, and measles were negatively enriched in response to Snhg15 suppression (Table 3). These results suggest that Snhg15 acts as a regulator of genes and pathways involved in antiviral responses, providing insight into how decreased Snhg15 expression may enhance VEEV replication.

**Table 3.** KEGG pathways shared at all time points after TC-83 infection in Snhg15 KD cells.

| Signaling pathways | Normalized Enrichment Score (NES) |  |  |
| --- | --- | --- | --- |
|  | 8hpi | 16hpi | 24hpi |
| TNF signaling pathway | -2.03 | -2.91 | -2.72 |
| Ribosome biogenesis in eukaryotes | -1.54 | -2.35 | -2.35 |
| NF-kappa B signaling pathway | -2 | -2.34 | -2.34 |
| IL-17 signaling pathway | -1.89 | -2.08 | -2.33 |
| Cytosolic DNA-sensing pathway | -1.78 | -1.87 | -2.15 |
| Coronavirus disease - COVID-19 | -1.45 | -1.62 | -2 |
| Epstein-Barr virus infection | -1.73 | -1.8 | -1.96 |
| Legionellosis | -1.51 | -1.81 | -1.92 |
| Rheumatoid arthritis | -1.71 | -1.65 | -1.88 |
| C-type lectin receptor signaling pathway | -1.5 | -1.64 | -1.87 |
| AGE-RAGE signaling pathway in diabetic complications | -1.37 | -1.94 | -1.8 |
| Chagas disease | -1.35 | -1.78 | -1.8 |
| NOD-like receptor signaling pathway | -1.68 | -1.73 | -1.73 |
| Human T-cell leukemia virus 1 infection | -1.5 | -1.58 | -1.71 |
| Chemokine signaling pathway | -1.36 | -1.65 | -1.71 |
| Kaposi sarcoma-associated herpesvirus infection | -1.56 | -1.68 | -1.7 |
| Lipid and atherosclerosis | -1.48 | -1.67 | -1.7 |
| Osteoclast differentiation | -1.51 | -1.55 | -1.62 |
| Hepatitis B | -1.46 | -1.9 | -1.62 |
| Hepatitis C | -1.48 | -1.69 | -1.6 |
| Viral protein interaction with cytokine and cytokine receptor | -1.9 | -1.55 | -1.59 |
| Toll-like receptor signaling pathway | -1.7 | -1.81 | -1.53 |
| Human cytomegalovirus infection | -1.39 | -1.44 | -1.52 |
| RIG-I-like receptor signaling pathway | -1.64 | -1.6 | -1.51 |
| Acute myeloid leukemia | -1.47 | -1.7 | -1.43 |
| Measles | -1.42 | -1.41 | -1.43 |
| Transcriptional misregulation in cancer | -1.49 | -1.42 | -1.38 |

Together, our data show for the first time that VEEV infection of primary target cells induces vastly different lncRNA profiles, dependent on infection with wild-type or attenuated VEEV. We identified lncRNAs modulating VEEV infection and highlighted lncRNA Snhg15 as a potential regulator of innate antiviral pathways during VEEV infection. The investigation of changes in the protein levels of genes suppressed at transcript level during Snhg15 KD in TC-83 infected cells will shed light on the mechanism Snhg15 uses to control VEEV infection.

## STAR ★ Methods

| REAGENT or RESOURCE | SOURCE | IDENTIFIER |
| --- | --- | --- |
| Virus Strains |  |  |
| VEEV Trinidad Donkey (VEEV IA/B) | BEI Resources | NR-332 |
| VEEV TC-83 (Vaccine strain) | BEI Resources | NR-63 |
| Experimental Models: Cells |  |  |
| Mouse C57 brain mixed primary astrocytes | Lonza | M-ASM-330 |
| Primary mouse cortical neurons | Gibco | A15586 |
| Vero E6 cells | ATCC | CRL-1586 |
| TaqMan Gene Expression Assays |  |  |
| A530013C23Rik | ThermoFisher Scientific | Mm01244073_m1 |
| 4930455J16Rik | Thermofisher scientific | Mm04278894_m1 |
| 3110039M20Rik | Thermofisher scientific | Mm01293535_m1 |
| Miat | Thermofisher scientific | Mm03455851_s1 |
| Mir155hg | Thermofisher Scientific | Mm01716204_m1 |
| Snhg15 | Thermofisher scientific | Mm01179930_m1 |
| Mir9-3hg | Thermofisher Scientific | Mm01319594_m1 |
| GM3734 | Thermofisher Scientific | ARNKU76 |
| GAPDH | Thermofisher scientific | Mm99999915_g1 |
| VEEV TC-83 | The sequence published by (Behnia et al., 2022) |  |
| siRNAs |  |  |
| Lincode Mouse Mir9-3hg(101694)-siRNA SMARTpool | Horizon discovery | R-160159-00-0005 |
| Lincode Mouse 3110039M20Rik(67293) siRNA-SMARTpool | Horizon discovery | R-173963-00-0005 |
| Lincode Mouse A530013C23Rik(329562) siRNA-SMARTpool | Horizon discovery | R-053806-00-0005 |
| Lincode Mouse Miat(330166) siRNA-SMARTpool | Horizon discovery | R-066744-00-0005 |
| Lincode Non-Targeting Pool -20nM | Horizon Discovery | D-001320-10-20 |
| Si-Snhg15 | The sequence published by (Du et al., 2018) |  |
| Plasmid |  |  |
| TC-83 plasmid | Kindly provided by Dr. Kyleen Khen-Hall |  |
| Reagents and commercial kits |  |  |
| ABM Basal Medium | Lonza | CC-3187 |
| AGM SingleQuots | Lonza | CC-4124 |
| Neurobasal Medium | Gibco | 21103 |
| GlutaMAX-I | Gibco | 35050 |
| B-27 | Gibco | 17504 |
| TransIT-X2 transfection reagent | Mirus | MIR6004 |
| Tri-reagent | Zymo Research | R2050-1-200 |
| Superscript II reverse transcriptase | Invitrogen | 18064014 |
| Direct-Zol RNA Microprep Kit | Zymo Research | R2060 |
| Qubit RNA High Sensitivity Assay Kit | Invitrogen | Q32855 |
| RiboGone Kit | Takara Bio | 634846 |
| Software |  |  |
| R version 4.2.2 | https://cran.rstudio.com/ |  |
| GraphPad Prism | https://www.graphpad.com/features |  |

### Resource availability

Further information and resources will be available through contacting the lead contact, Dr. Steven B. Bradfute.

### Material availability

This research did not generate novel reagents.

### Data and code availability

RNA sequencing files will be made available upon manuscript acceptance in a public database. Additional information required for reanalyzing the data reported in this paper is available upon request to the lead contact, Dr. Steven B. Bradfute.

## Acknowledgements

Research reported in this publication was supported by the Defense Threat Reduction Agency (HDTRA-1-20-1-0015 (SBB) and an Institutional Development Award (IDeA) from the National Institute of General Medical Sciences of the National Institutes of Health under grant number P20GM103451. The content of the information does not necessarily reflect the position or the policy of the federal government, and no official endorsement should be inferred.

## Material and Methods

### Cell Culture

Mouse C57 brain mixed primary astrocytes (Lonza, M-ASM-330) were grown in ABM Basal Medium (Lonza, CC-3187) supplemented with AGM SingleQuots (Lonza, CC-4124). At passage five, cells were seeded into 6-well plates (5 × 10^5 cells/well) maintained in supplemented ABM Basal Medium at 37 °C and 5% CO2 for 24 hours. Primary mouse cortical neurons (Gibco, A15586) were seeded into 6-well plates (1 × 10^6 cells/well) four days before infection. The plates were precoated with poly-D-lysine (25 μg/mL) and cells were maintained in Neurobasal Medium (Gibco, 21103) supplemented with 0.5 mM GlutaMAX-I (Gibco, 35050) and 2% (v/v) B-27 (Gibco, 17504) at 37°C and 5% CO2 until infection. Vero E6 cells (ATCC, CRL-1586) were cultured in 6-well plates (0.3 × 10^6 cells/well) and maintained at 37°C and 5% CO2 in Dulbecco’s Modified Eagle Medium (DMEM) supplemented with 10% fetal bovine serum (FBS) and 1% Pen Strep Glutamine (100×) (Gibco, 10378016).

### Viruses

Wild type VEEV TrD and the live attenuated vaccine VEEV TC-83 strains obtained from the Biodefense and Emerging Infections Research Resources Repository (BEI Resources, NR-332 and NR-63). Both viral stocks were expanded in Vero E6 cells and quantified using plaque assays. All TrD work was conducted in an approved select agent BSL3 facility, and RNA was inactivated prior to removal for sequencing as described below.

### RNA sequencing of VEEV infected cells

Primary mouse astrocytes at passage five and primary mouse neurons were cultured in 6-well plates as explained in the cell culture section. Infections were performed 24 hours after seeding for primary mouse astrocytes (three biological replicates) and four days post-seeding for primary mouse neurons (three biological replicates were infected with TC-83 and two with TrD). On the day of infection, the cells were infected with either VEEV TC-83 or VEEV TrD at MOI 5 for one hour. After one hour, the viral inoculum was replaced with fresh pre-warmed cell culture medium, and the cells were incubated at 37°C and 5% CO2. Total RNA was extracted from uninfected cells and infected cells at 16- and 24-hours post-infection using the Direct-Zol RNA Microprep Kit (Zymo Research, R2060) according to the manufacturer’s protocol. Total RNA was quantified using the Qubit RNA High Sensitivity Assay Kit (Invitrogen, Q32855) on a Qubit4, following the manufacturer’s protocol. The total RNA extracted from infected and uninfected cells was fragmented as described previously (Behnia et al., 2022). Ribosomal RNA was depleted using the RiboGone Kit (Takara Bio, Cat. No. 634846) according to the manufacturer’s instructions. The RNA was prepared for sequencing using the SMARTer Universal Low Input RNA Kit for Sequencing (Takara Bio) following the manufacturer’s instructions. Samples were then prepared using the Ion Plus Fragment Library Kit (Ion Torrent) at the UNM ATG core facility, starting at the Ligation and Nick Repair step, with the only modification being the use of 80% ethanol instead of 70% ethanol for bead washes. Samples were loaded onto the Ion 540 Chip (Ion Torrent) and run on the Ion S5 XL Machine. These methods were previously published by Brown et al. and Brayer et al. (Brayer et al., 2016; Brown et al., 2017). The RNA-seq reads were aligned to the mouse reference genome (MM10) using the STAR aligner. Differential gene expression analysis was performed using the DESeq2 library in R version 4.2.2. KEGG pathway analyses were conducted in R version 4.2.2 using the clusterProfiler, enrichplot, ggplot2, and Tidyverse libraries. The KEGG pathway analysis was performed using the R codes published at https://learn.gencore.bio.nyu.edu/rna-seq-analysis/gene-set-enrichment-analysis/

### Validation of lncRNA Expression

Five biological replicates of primary mouse astrocytes at passage five were cultured in 6-well plates (5 × 10^5 cells/well) and maintained in ABM Basal Medium (Lonza, CC-3187) supplemented with AGM SingleQuots (Lonza, CC-4124) at 37°C and 5% CO2 for 24 hours. After 24 hours, cells were incubated with VEEV TC-83 (MOI 5) for one hour or left uninfected as controls. The viral inoculum was replaced with cell culture medium one hour after infection, and cells were incubated at 37°C and 5% CO2. Infected and uninfected cells were lysed using Tri-reagent (Zymo Research, R2050-1-200) at 16- and 24-hour post-infection. Total RNA was extracted from the cell lysates using the Direct-Zol RNA Microprep Kit (Zymo Research, R2060) according to the manufacturer’s protocol. The extracted RNAs were stored at -80°C for further RT-qPCR analysis.

### RT-qPCR - Validation of lncRNA expression

500 ng of total RNA extracted from uninfected or VEEV TC-83 (MOI 5) infected primary mouse astrocytes at 16- and 24-hours post-infection were subjected to cDNA synthesis using Superscript II reverse transcriptase (Invitrogen, 18064014) according to the manufacturer’s protocol. The amount of RNA was back-calculated in cDNA to use 25ng of RNA/reaction. TaqMan gene expression assays, including Mm01244073_m1 (A530013C23Rik), Mm04278894_m1 (4930455J16Rik), Mm01293535_m1 (3110039M20Rik), Mm03455851_s1 (Miat), Mm01716204_m1 (Mir155hg), Mm01179930_m1 (Snhg15), Mm01319594_m1 (Mir9-3hg), ARNKU76 (GM3734), and Mm99999915_g1 (GAPDH) were purchased from ThermoFisher Scientific to measure the expression levels of lncRNAs and GAPDH. The qPCR was performed on a QuantStudio 5 Real-Time PCR Machine (Applied Biosystems, USA), with two wells tested per sample. The expression levels of lncRNAs in each sample were measured relative to GAPDH and normalized to the expression of lncRNAs in uninfected cells using the ΔΔCt method. Five biological replicates of primary mouse astrocytes were used in this assay. Graphs were generated using GraphPad Prism (Version 10), and one-way ANOVA followed by multiple comparisons was used to assess statistical changes in the expression of lncRNAs.

### siRNA Screening

Three biological replicates of mouse C57 primary mouse astrocytes (Lonza, M-ASM-330) at passage five cultured into 6-well plates (5 × 10^5^ cells/ well) and maintained in ABM Basal Medium (Lonza, CC-3187) supplemented with AGM SingleQuots (Lonza, CC-4124) at 37°C and 5% CO_2_. The siRNA SMARTpools purchased from for this assay are as follows: Lincode Mouse Mir9-3hg (101694) siRNA-SMARTpool (Horizon Discovery, R-160159-00-0005), Lincode Mouse 3110039M20Rik (67293) siRNA-SMARTpool (Horizon Discovery, R-173963-00-0005), Lincode Mouse A530013C23Rik (329562) siRNA-SMARTpool (Horizon discovery, R-053806-00-0005), Lincode Mouse Miat (330166) siRNA-SMARTpool (Horizon Discovery, R-066744-00-0005) and Lincode Non-Targeting Pool (Horizon Discovery, D-001320-10-20). The sequence of the siRNA used to suppress Snhg15 can be found in a paper by (Du et al., 2018). Twenty-four hours post-seeding cells were transfected with 50nM of siRNA targeting lncRNA or a pool of nontargeting siRNAs (control) using TransIT-X2 transfection reagent (Mirus, MIR6004) using the manufacturer’s protocol, with the only modification being the use of ½ of recommended transfection reagent/rxn. Each siRNApool was transfected to two wells, except for siRNA used against Snhg15 that transfected to one well. Twenty-four hours after transfection cells were infected with VEEV TC-83 (MOI 5) for one hour or left uninfected (control), with the exception of cells transfected with siRNA against Snhg15 that were infected with TC-83 at MOI 0.5. The viral inoculum replaced with cell culture medium after one hour and cells incubated at 37°C and 5% CO_2_ for 24 hours. At 24h post-infection cells lysed using Tri-reagent (Zymo-research, R2050-1-200) and total RNA extracted from the cell lysates using Direct-Zol RNA Microprep Kit (Zymo Research, R2060) according to the manufacturer protocol. The total RNAs stored at -80C for later RT-qPCR analysis.

### RT-qPCR - Validation of lncRNA suppression

To measure the expression levels of lncRNAs after lncRNA suppression. Total RNA extracted from cells treated with a pool of four nontargeting siRNA (control for baseline lncRNA expression), treated with siRNA targeting lncRNA followed by infection with VEEV TC-83 (MOI 5 or 0.5), and transfected with a pool of four nontargeting siRNAs followed by infection with VEEV TC-83 (MOI 5 or 0.5)were subjected to cDNA synthesis using Superscript II reverse transcriptase (Invitrogen, 18064014) according to the manufacturer protocol. The volume of cDNA containing 25ng total RNA was used in each reaction.. TaqMan gene expression assays, Mm01244073_m1(A530013C23Rik), Mm01293535_m1(3110039M20Rik), Mm01179930_m1 (Snhg15), Mm01319594_m1 (Mir9-3hg), Mm03455851_s1 (Miat) and Mm99999915_g1 (GAPDH) purchased from ThermoFisher Scientific to measure the expression levels of lncRNAs and GAPDH. The qPCR was performed in QuantStudio 5 Realtime PCR Machine (Applied Biosystems, USA). The expression of each lncRNA tested in four qPCR wells, with the exception of Snhg15 expression that was tested in three wells.. The expression level of lncRNA in each sample calculated relative to GAPDH and normalized to the expression of lncRNA in primary mouse astrocytes transfected with a pool of non-targeting siRNA using ΔΔCt method. Three biological replicates of primary mouse astrocytes were used in this assay.

### Plaque assays

To measure the potential effect of lncRNA suppression on the viral titer, Vero E6 cells were seeded into 6-well plates (0.5 × 10^6^ cells/well) 24h prior to infection. Twenty-four hours post-seeding, the cells were incubated with ten-fold serial dilutions of supernatants from lncRNA suppressed primary mouse astrocytes that are infected with VEEV TC-83 (MOI 5), primary mouse astrocytes transfected with a pool of four non-targeting siRNAs (negative control) or primary mouse astrocytes transfected with a pool of four non-targeting siRNAs followed by infection with VEEV TC-83 (MOI 5) (positive control) for one hour at 37°C with rocking at 30 min. The supernatant used for plaque assay were collected at 24hpi. Dilutions 10^−1^ through 10^−9^ of the supernatants used in plaque assay. After the incubation period, the inoculum was replaced with 1ml of 1% agarose in Modified Eagle Medium supplemented with 2.5% FBS. The plates were incubated at 37°C and 5% CO_2_ for 48h. To fix the cells, 1 mL of 4% formaldehyde was added to the agar overlay and the plates incubated at 4°C overnight. After the incubation period, the agar overlay was removed, and the fixed monolayer was stained with 0.8% crystal violet solution. Plaques were quantified and virus concentrations were recorded as PFU/mL. Three wells per dilution were tested. Three biological replicates were used in this assay.

### RNA sequencing of Snhg15 knockdown cells

Three biological replicates of primary mouse astrocytes (Lonza, M-ASM-330) at passage five were cultured into 6-well plates and maintained in astrocytes growth medium at 37°C and 5% CO_2_ for 24 hours. At 24-hour post-seeding, cells were transfected with either siRNA targeting Snhg15 or a pool of four non-targeting siRNAs (Control) using TransIT-X2 transfection reagent (Mirus, MIR6004) according to the manufacturer’s protocol with modification of using ½ of suggested transfection reagent. The transfected cells incubated at 37°C and 5% CO_2_ for 24 hours. 24h after transfection, cells were incubated with VEEV TC-83 (MOI 5) for one hour with rocking at 30 minutes. After one-hour, viral inoculum replaced with cell culture medium, and cells incubated at 37°C and 5% CO_2_ for another 24 hours. At 24h post-infection cells were lysed using Tri-reagent (Zymo-research, R2050-1-200) and total RNA extracted from the cell lysates using Direct-Zol RNA Microprep Kit (Zymo Research, R2060) according to the manufacturer protocol. Total RNA samples were sent to the University of Colorado-Anschutz sequencing center for RNA sequencing. After the ribosomal RNA depletion, the RNA sequencing ran on the NovaSeqX Machine. The paired end sequencing was done with the read depth of 50 million per sample. The obtained reads were aligned to mouse reference genome (MM10) using STAR aligner and differential gene expression analysis was done in R version 4.2.2. using DESeq2 library. KEGG pathway analyses were performed in R version 4.2.2. using clusterProfiler, enrichplot, ggplot2, and tidyverse libraries and using R codes published at https://learn.gencore.bio.nyu.edu/rna-seq-analysis/gene-set-enrichment-analysis/. Venn diagram generated in R using version 4.2.2 and using VennDiagram and grid libraries.

## Supplementary Figures

**Supplementary Figure 1.**
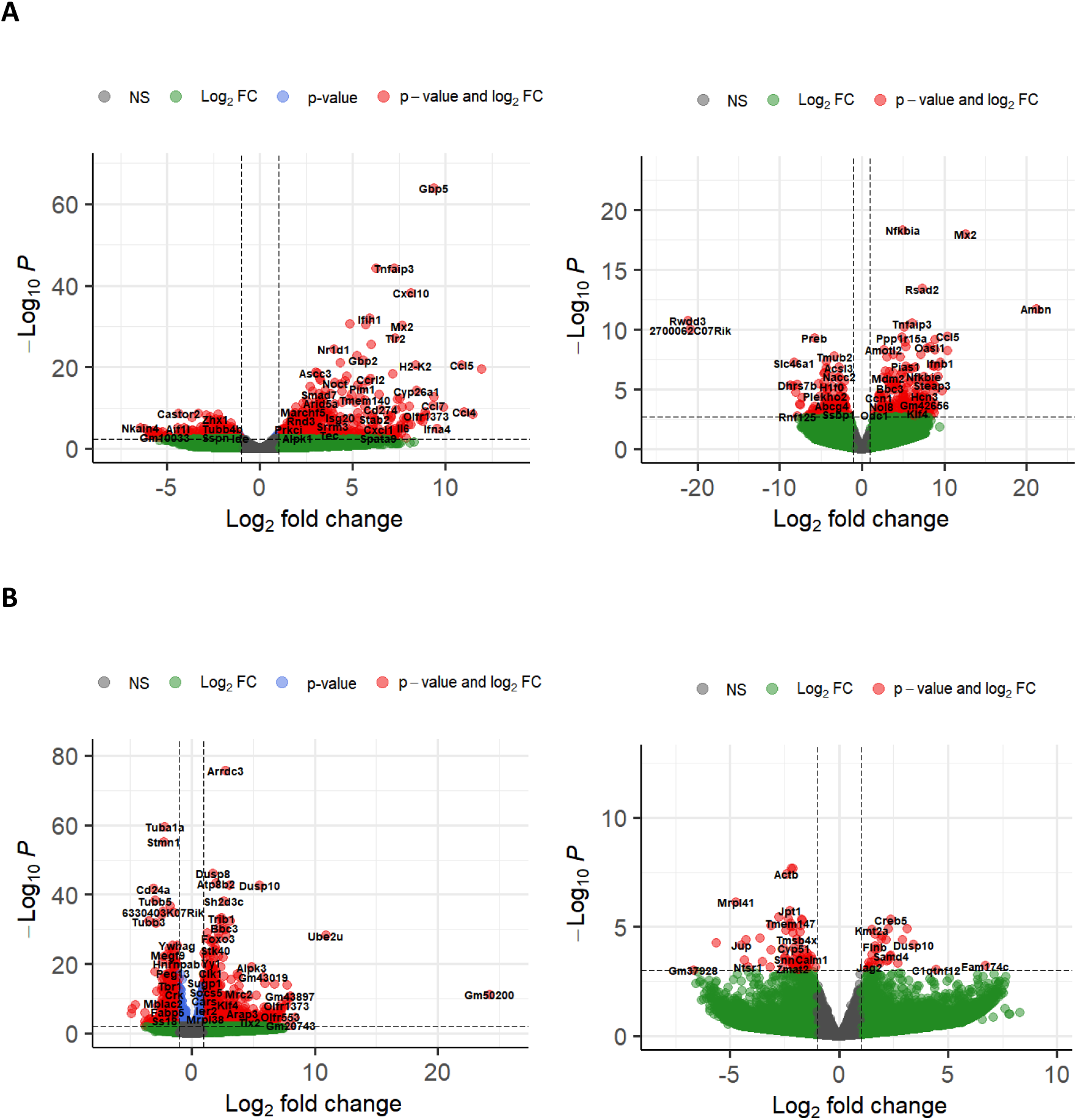
The host cellular response to VEEV varies depending on the VEEV strain and cell type. Volcano plots show DEG in primary mouse astrocytes **(A)**, primary mouse neurons **(B)**, infected with eighter VEEV TC-83 (Left) or VEEV TrD (Right) for 24hours. p-value threshold adjusted to p.Adj value = 0.05

**Supplementary Figure 2.**
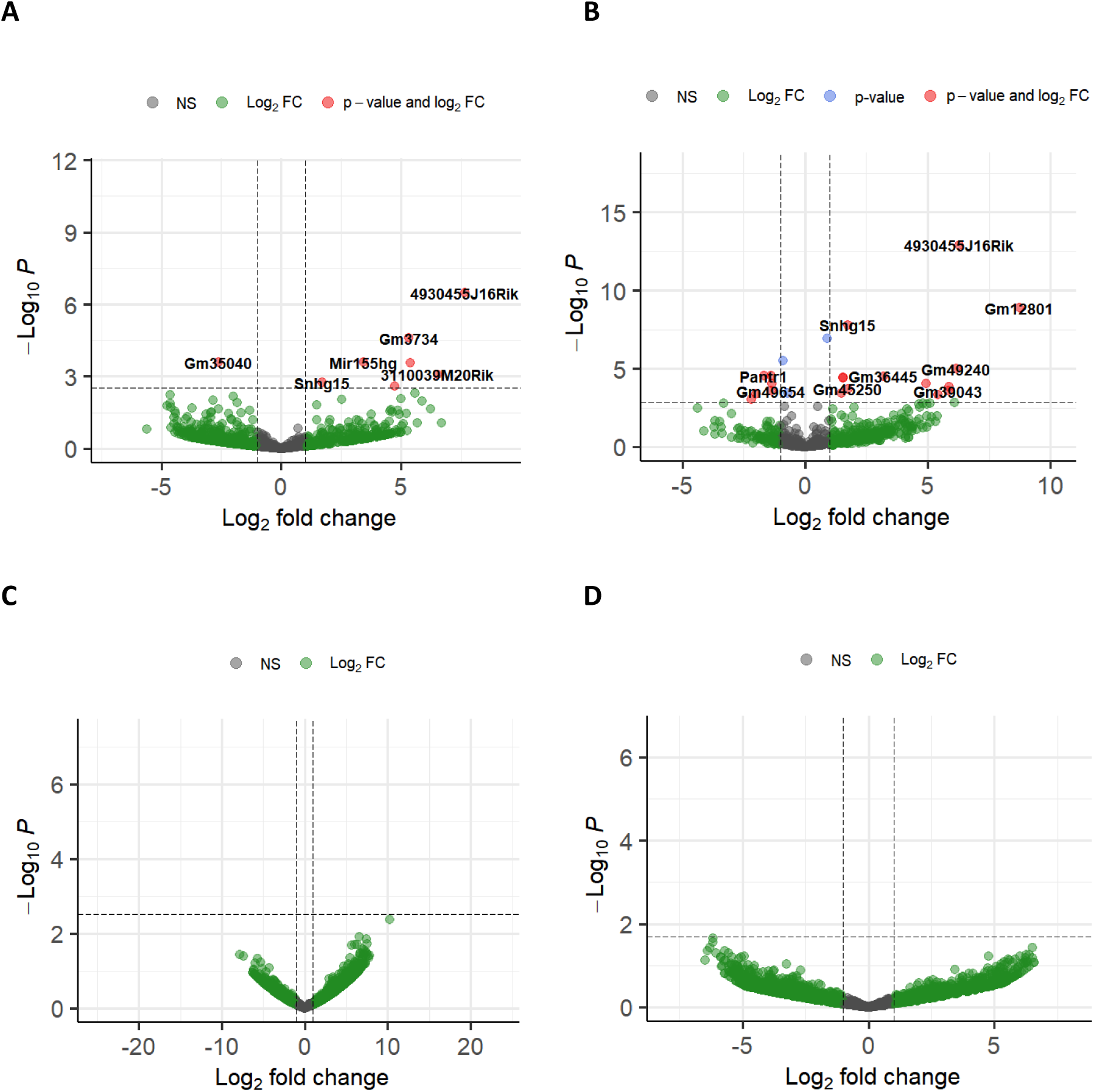
The host cellular lncRNA response to VEEV strains at 16hpi. **A-B)** Volcano plots show DE-lncRNAs in primary mouse astrocytes **(A)** and primary mouse neurons **(B)** infected with VEEV TC-83. **C-D)** Volcano plots show DE-lncRNAs in primary mouse astrocytes **(C)** and primary mouse neurons **(D)** infected with VEEV TrD. All the plots show the results from RNA-seq at 16h.p.i. The p-value threshold adjusted to show p.Adj value = 0.05.

## References

Aguilar, P. V., Estrada-Franco, J. G., Navarro-Lopez, R., Ferro, C., Haddow, A. D., & Weaver, S. C. (2011). Endemic Venezuelan equine encephalitis in the Americas: Hidden under the dengue umbrella. Future Virology, 6(6), 721–740. 10.2217/FVL.11.5

Badu, P., & Pager, C. T. (2023). Activation of ATF3 via the Integrated Stress Response Pathway Regulates Innate Immune and Autophagy Processes to Restrict Zika Virus. bioRxiv, 2023.07.26.550716. 10.1101/2023.07.26.550716

Behnia, M., Baer, A., Skidmore, A. M., Lehman, C. W., Bracci, N., Kehn-Hall, K., & Bradfute, S. B. (2022). Inactivation of Venezuelan Equine Encephalitis Virus Genome Using Two Methods. Viruses, 14(2), Article 2. 10.3390/v14020272

Bhalla, N., Sun, C., Metthew Lam, L. K., Gardner, C. L., Ryman, K. D., & Klimstra, W. B. (2016). Host translation shutoff mediated by non-structural protein 2 is a critical factor in the antiviral state resistance of Venezuelan equine encephalitis virus. Virology, 496, 147–165. 10.1016/j.virol.2016.06.005

Brayer, K. J., Frerich, C. A., Kang, H., & Ness, S. A. (2016). Recurrent Fusions in MYB and MYBL1 Define a Common, Transcription Factor-Driven Oncogenic Pathway in Salivary Gland Adenoid Cystic Carcinoma. Cancer Discovery, 6(2), 176–187. 10.1158/2159-8290.CD-15-0859

Brown, R. B., Madrid, N. J., Suzuki, H., & Ness, S. A. (2017). Optimized approach for Ion Proton RNA sequencing reveals details of RNA splicing and editing features of the transcriptome. PLOS ONE, 12(5), e0176675. 10.1371/journal.pone.0176675

Chen, S., Huang, X., Xie, Q., Liu, Q., & Zhu, H. (2022). The Role of Long Noncoding RNA BST2-2 in the Innate Immune Response to Viral Infection. Journal of Virology, 96(7). 10.1128/jvi.00207-22

De la Fuente-Hernandez, M. A., Sarabia-Sanchez, M. A., Melendez-Zajgla, J., & Maldonado-Lagunas, V. (2022). lncRNAs in mesenchymal differentiation. American Journal of Physiology. Cell Physiology, 322(3), C421–C460. 10.1152/ajpcell.00364.2021

Du, Y., Kong, C., Zhu, Y., Yu, M., Li, Z., Bi, J., Li, Z., Liu, X., Zhang, Z., & Yu, X. (2018). Knockdown of SNHG15 suppresses renal cell carcinoma proliferation and EMT by regulating the NF-κB signaling pathway. International Journal of Oncology, 53(1), 384–394. 10.3892/ijo.2018.4395

Eaton, B. (2021). Cellular Immune Evasion Mechanisms of New World Encephalitic Alphaviruses [Ph.D., The George Washington University]. https://www.proquest.com/docview/2572536418/abstract/1D3341EC84474BDDPQ/1

Forrester, N. L., Wertheim, J. O., Dugan, V. G., Auguste, A. J., Lin, D., Adams, A. P., Chen, R., Gorchakov, R., Leal, G., Estrada-Franco, J. G., Pandya, J., Halpin, R. A., Hari, K., Jain, R., Stockwell, T. B., Das, S. R., Wentworth, D. E., Smith, M. D., Kosakovsky Pond, S. L., & Weaver, S. C. (2017). Evolution and spread of Venezuelan equine encephalitis complex alphavirus in the Americas. PLoS Neglected Tropical Diseases, 11(8), e0005693. 10.1371/journal.pntd.0005693

Garmashova, N., Atasheva, S., Kang, W., Weaver, S. C., Frolova, E., & Frolov, I. (2007). Analysis of Venezuelan Equine Encephalitis Virus Capsid Protein Function in the Inhibition of Cellular Transcription. Journal of Virology, 81(24), 13552–13565. 10.1128/JVI.01576-07

Ginn, L., La Montagna, M., Wu, Q., & Shi, L. (2021). Diverse roles of long non-coding RNAs in viral diseases. Reviews in Medical Virology, 31(4). 10.1002/rmv.2198

Grieder, F. B., & Vogel, S. N. (1999). Role of Interferon and Interferon Regulatory Factors in Early Protection against Venezuelan Equine Encephalitis Virus Infection. Virology, 257(1), 106–118. 10.1006/viro.1999.9662

Gupta, P., Sharma, A., Han, J., Yang, A., Bhomia, M., Knollmann-Ritschel, B., Puri, R. K., & Maheshwari, R. K. (2017). Differential host gene responses from infection with neurovirulent and partially-neurovirulent strains of Venezuelan equine encephalitis virus. BMC Infectious Diseases, 17. 10.1186/s12879-017-2355-3

Hai, T., Wolfgang, C. D., Marsee, D. K., Allen, A. E., & Sivaprasad, U. (2018). ATF3 and Stress Responses. Gene Expression, 7(4-5–6), 321–335.

Kaczynski, T. J., Husami, N. J., Au, E. D., & Farkas, M. H. (2023). Dysregulation of a lncRNA within the TNFRSF10A locus activates cell death pathways. Cell Death Discovery, 9(1), 242. 10.1038/s41420-023-01544-5

Korbecki, J., Barczak, K., Gutowska, I., Chlubek, D., & Baranowska-Bosiacka, I. (2022). CXCL1: Gene, Promoter, Regulation of Expression, mRNA Stability, Regulation of Activity in the Intercellular Space. International Journal of Molecular Sciences, 23(2), Article 2. 10.3390/ijms23020792

Koterski, J., Twenhafel, N., Porter, A., Reed, D. S., Martino-Catt, S., Sobral, B., Crasta, O., Downey, T., & DaSilva, L. (2007). Gene expression profiling of nonhuman primates exposed to aerosolized Venezuelan equine encephalitis virus. FEMS Immunology & Medical Microbiology, 51(3), 462–472. 10.1111/j.1574-695X.2007.00319.x

Lai, K. M., Goh, B. H., & Lee, W. L. (2020). Attenuating influenza a virus infection by heparin binding EGF-like growth factor. *Growth Factors (Chur*, Switzerland), 38(3–4), 167–176. 10.1080/08977194.2021.1895144

Lundberg, L., Carey, B., & Kehn-Hall, K. (2017). Venezuelan Equine Encephalitis Virus Capsid—The Clever Caper. Viruses, 9(10). 10.3390/v9100279

Mattick, J. S., Amaral, P. P., Carninci, P., Carpenter, S., Chang, H. Y., Chen, L.-L., Chen, R., Dean, C., Dinger, M. E., Fitzgerald, K. A., Gingeras, T. R., Guttman, M., Hirose, T., Huarte, M., Johnson, R., Kanduri, C., Kapranov, P., Lawrence, J. B., Lee, J. T., … Wu, M. (2023). Long non-coding RNAs: Definitions, functions, challenges and recommendations. Nature Reviews Molecular Cell Biology, 24(6), 430–447. 10.1038/s41580-022-00566-8

Nair, S., Poddar, S., Shimak, R. M., & Diamond, M. S. (2017). Interferon Regulatory Factor 1 Protects against Chikungunya Virus-Induced Immunopathology by Restricting Infection in Muscle Cells. Journal of Virology, 91(22), e01419–17. 10.1128/JVI.01419-17

Okamura, M., Shizu, R., Abe, T., Kodama, S., Hosaka, T., Sasaki, T., & Yoshinari, K. (2020). PXR Functionally Interacts with NF-κB and AP-1 to Downregulate the Inflammation-Induced Expression of Chemokine CXCL2 in Mice. Cells, 9(10), Article 10. 10.3390/cells9102296

Panda, D., Gjinaj, E., Bachu, M., Squire, E., Novatt, H., Ozato, K., & Rabin, R. L. (2019). IRF1 Maintains Optimal Constitutive Expression of Antiviral Genes and Regulates the Early Antiviral Response. Frontiers in Immunology, 10, 1019. 10.3389/fimmu.2019.01019

Penkala, I., Wang, J., Syrett, C. M., Goetzl, L., López, C. B., & Anguera, M. C. (2016). lncRHOXF1, a Long Noncoding RNA from the X Chromosome That Suppresses Viral Response Genes during Development of the Early Human Placenta. Molecular and Cellular Biology, 36(12), 1764–1775. 10.1128/MCB.01098-15

Pittman, P. R., Makuch, R. S., Mangiafico, J. A., Cannon, T. L., Gibbs, P. H., & Peters, C. J. (1996). Long-term duration of detectable neutralizing antibodies after administration of live-attenuated VEE vaccine and following booster vaccination with inactivated VEE vaccine. Vaccine, 14(4), 337–343. 10.1016/0264-410x(95)00168-z

Rusnak, J. M., Dupuy, L. C., Niemuth, N. A., Glenn, A. M., & Ward, L. A. (2018). Comparison of Aerosol- and Percutaneous-acquired Venezuelan Equine Encephalitis in Humans and Nonhuman Primates for Suitability in Predicting Clinical Efficacy under the Animal Rule. Comparative Medicine, 68(5), 380–395. 10.30802/AALAS-CM-18-000027

Saha, I., Jaiswal, H., Mishra, R., Nel, H. J., Schreuder, J., Kaushik, M., Singh Chauhan, K., Singh Rawat, B., Thomas, R., Naik, S., Kumar, H., & Tailor, P. (2020). RelB suppresses type I Interferon signaling in dendritic cells. Cellular Immunology, 349, 104043. 10.1016/j.cellimm.2020.104043

Schmitz, S. U., Grote, P., & Herrmann, B. G. (2016). Mechanisms of long noncoding RNA function in development and disease. Cellular and Molecular Life Sciences, 73, 2491–2509. 10.1007/s00018-016-2174-5

Sharma, A., Bhattacharya, B., Puri, R. K., & Maheshwari, R. K. (2008). Venezuelan equine encephalitis virus infection causes modulation of inflammatory and immune response genes in mouse brain. BMC Genomics, 9, 289. 10.1186/1471-2164-9-289

Sharma, A., Bhomia, M., Honnold, S. P., & Maheshwari, R. K. (2011). Role of adhesion molecules and inflammation in Venezuelan equine encephalitis virus infected mouse brain. Virology Journal, 8(1), 197. 10.1186/1743-422X-8-197

Sharma, A., & Knollmann-Ritschel, B. (2019). Current Understanding of the Molecular Basis of Venezuelan Equine Encephalitis Virus Pathogenesis and Vaccine Development. Viruses, 11(2). 10.3390/v11020164

Sharma, A., & Maheshwari, R. K. (2009). Oligonucleotide array analysis of Toll-like receptors and associated signalling genes in Venezuelan equine encephalitis virus-infected mouse brain. The Journal of General Virology, 90(Pt 8), 1836–1847. 10.1099/vir.0.010280-0

Shuai, Y., Ma, Z., Lu, J., & Feng, J. (2019). LncRNA SNHG15: A new budding star in human cancers. Cell Proliferation, 53(1), e12716. 10.1111/cpr.12716

Statello, L., Guo, C.-J., Chen, L.-L., & Huarte, M. (2021). Gene regulation by long non-coding RNAs and its biological functions. Nature Reviews Molecular Cell Biology, 22(2), Article 2. 10.1038/s41580-020-00315-9

Sun, H., Li, S., Xu, Z., Liu, C., Gong, P., Deng, Q., & Yan, F. (2022). SNHG15 is a negative regulator of inflammation by mediating TRAF2 ubiquitination in stroke-induced immunosuppression. Journal of Neuroinflammation, 19, 1. 10.1186/s12974-021-02372-z

Valadkhan, S., & Gunawardane, L. S. (2016). lncRNA-mediated regulation of the interferon response. Virus Research, 212, 127–136. 10.1016/j.virusres.2015.09.023

Valerol, N., Bonilla, E., Espina, L. M., Maldonado, M., Montero, E., Añez, F., Levy, A., Bermudez, J., Meleán, E., & Nery, A. (2008). [Increase of interleukin-1 beta, gamma interferon and tumor necrosis factor alpha in serum and brain of mice infected with the Venezuelan Equine Encephalitis virus]. Investigacion Clinica, 49(4), 457–467.

Vierbuchen, T., & Fitzgerald, K. A. (2021). Long non-coding RNAs in antiviral immunity. Seminars in Cell & Developmental Biology, 111, 126–134. 10.1016/j.semcdb.2020.06.009

Watts, D. M., Callahan, J., Rossi, C., Oberste, M. S., Roehrig, J. T., Wooster, M. T., Smith, J. F., Cropp, C. B., Gentrau, E. M., Karabatsos, N., Gübler, D., & Hayes, C. G. (1998). Venezuelan equine encephalitis febrile cases among humans in the Peruvian Amazon River region. The American Journal of Tropical Medicine and Hygiene, 58(1), 35–40. 10.4269/ajtmh.1998.58.35

Wickremasinghe, M. I., Thomas, L. H., O’Kane, C. M., Uddin, J., & Friedland, J. S. (2004). Transcriptional Mechanisms Regulating Alveolar Epithelial Cell-specific CCL5 Secretion in Pulmonary Tuberculosis*. Journal of Biological Chemistry, 279(26), 27199–27210. 10.1074/jbc.M403107200

Xu, J., Wang, P., Li, Z., Li, Z., Han, D., Wen, M., Zhao, Q., Zhang, L., Ma, Y., Liu, W., Jiang, M., Zhang, X., & Cao, X. (2021). IRF3-binding lncRNA-ISIR strengthens interferon production in viral infection and autoinflammation. Cell Reports, 37(5), 109926. 10.1016/j.celrep.2021.109926

Yao, Z., Zanini, F., Kumar, S., Karim, M., Saul, S., Bhalla, N., Panpradist, N., Muniz, A., Narayanan, A., Quake, S. R., & Einav, S. (2021). The transcriptional landscape of Venezuelan equine encephalitis virus (TC-83) infection. PLoS Neglected Tropical Diseases, 15(3), e0009306. 10.1371/journal.pntd.0009306

Zhang, Y., Chi, X., Hu, J., Wang, S., Zhao, S., Mao, Y., Peng, B., Chen, J., & Wang, S. (2023). LncRNA LINC02574 Inhibits Influenza A Virus Replication by Positively Regulating the Innate Immune Response. International Journal of Molecular Sciences, 24(8), 7248. 10.3390/ijms24087248

Zhou, F., Wan, Q., Chen, Y., Chen, S., & He, M.-L. (2021). PIM1 kinase facilitates Zika virus replication by suppressing host cells’ natural immunity. Signal Transduction and Targeted Therapy, 6(1), 207. 10.1038/s41392-021-00539-x

Zhou, H., Tang, Y.-D., & Zheng, C. (2022). Revisiting IRF1-mediated antiviral innate immunity. Cytokine & Growth Factor Reviews, 64, 1–6. 10.1016/j.cytogfr.2022.01.004

